# *Clostridioides difficile* bile salt hydrolase activity has substrate specificity and affects biofilm formation

**DOI:** 10.1101/2022.07.19.500743

**Authors:** Andrea Martinez Aguirre, Joseph A. Sorg

## Abstract

The *Clostridioides difficile* pathogen is responsible for nosocomial infections. Germination is an essential step for the establishment of *C. difficile* infection (CDI) because toxins that are secreted by vegetative cells are responsible for the symptoms of CDI. Germination can be stimulated by the combinatorial actions of certain amino acids and either conjugated or deconjugated cholic acid-derived bile salts. During synthesis in the liver, cholic acid- and chenodeoxycholic acid-class bile salts are conjugated with either taurine or glycine at the C24 carboxyl. During GI transit, these conjugated bile salts are deconjugated by microbes that express bile salt hydrolases (BSHs). Here, we surprisingly find that several *C. difficile* strains have BSH activity. We observed this activity in both *C. difficile* vegetative cells and in spores and that the observed BSH activity was specific to taurine-derived bile salts. Additionally, we find that this BSH activity can produce cholate for metabolic conversion to deoxycholate by *C. scindens*. The *C. scindens*-produced deoxycholate signals to *C. difficile* to initiate biofilm formation. Our results show that *C. difficile* BSH activity has the potential to influence the interactions between microbes and this could extend to the GI setting.

**Importance:** Both primary and secondary bile salts are well-established to impact *C. difficile* spore germination and vegetative growth. Here, we find that *C. difficile* vegetative cells, and spores, have bile salt hydrolase activity that is specific to taurine-derived bile salts. When grown in co-culture with the secondary bile salt-producing bacterium, *C. scindens*, we find that *C. difficile*-mediated deconjugation of taurocholate, ‘feeds’ *C. scindens* cholate. *C. scindens* 7α-dehydroxylates cholate to deoxycholate. The *C. scindens-*produced deoxycholate then stimulates biofilm formation by *C. difficile* cells. Thus, this suggests that the bile salt hydrolase activity expressed by several *C. difficile* strains could be responsible for modulating *in vivo* biofilm formation and maintenance in a host.

## Introduction

*C. difficile* is a Gram-positive, spore-forming, pathogenic bacterium that is considered the main cause of antibiotic associated diarrhea. The infectious agent of *C. difficile* is the spore form since dormant spores can persist in the environment for long periods of time and are resistant to commonly used disinfectants (1, 2). Nevertheless, CDI symptoms are a result of the TcdA and TcdB toxins that are secreted by *C. difficile* vegetative cells (2-4). Thus, germination of the dormant *C. difficile* spores to the growing vegetative cells is essential for disease development. In *C. difficile* the germination process can be triggered by certain host-derived bile salts and certain amino acids (5-11).

Bile salts are cholesterol-based molecules that are synthesized in the liver and, in humans, consist of the cholic acid (CA) and chenodeoxycholic acid (CDCA) base structures (Figure 1). Subsequently these base molecules are further modified by the addition of either taurine or glycine at C-24 [yielding taurocholic acid (TA) / taurochenodeoxycholic acid (TCDCA) and glycocholic acid (GCA) / glycochenodeoxycholic acid (GCDCA), respectively (Figure 1)]. These conjugated bile salts are then secreted into the intestines where they aid in the absorption of fats and cholesterol (12, 13). Both the conjugated and deconjugated forms of the cholic acid-class bile salts promote *C. difficile* spore germination, whereas chenodeoxycholic acid, and its derivatives, inhibit germination (5, 9-11, 14-19). Through an enterohepatic recirculation process, the majority of the intestinal bile salts are reabsorbed and sent back to the liver for other rounds of digestion. Bile salts that do not undergo enterohepatic recirculation can be deconjugated by bile salt hydrolases (BSHs), which remove the conjugated amino acid at the C-24 position (20-22). Deconjugated primary bile salts can then undergo additional biotransformations, such as epimerization, oxidation, 7α/β-dehydroxylation and dehydrogenation (20, 23). The 7α-dehydroxylation process, performed by a small subset of gut microbes, generates the secondary bile salts deoxycholic acid (DCA) from CA and lithocholic acid (LCA) from CDCA (Figure 1). Secondary bile salts strongly correlate with, but may not result in, an environment that resists *C. difficile* colonization (24-28). Because bile salt deconjugation is a requirement for downstream bile salt modifications, the role of BSH enzymes has been studied due to their effects on bile salt regulation and potential effects on lowering cholesterol levels, management of obesity, and the effects in gut inflammatory disorders (*e*.*g*., inflammatory bowel diseases and type 2 diabetes). (29-33).

**Figure 1.**
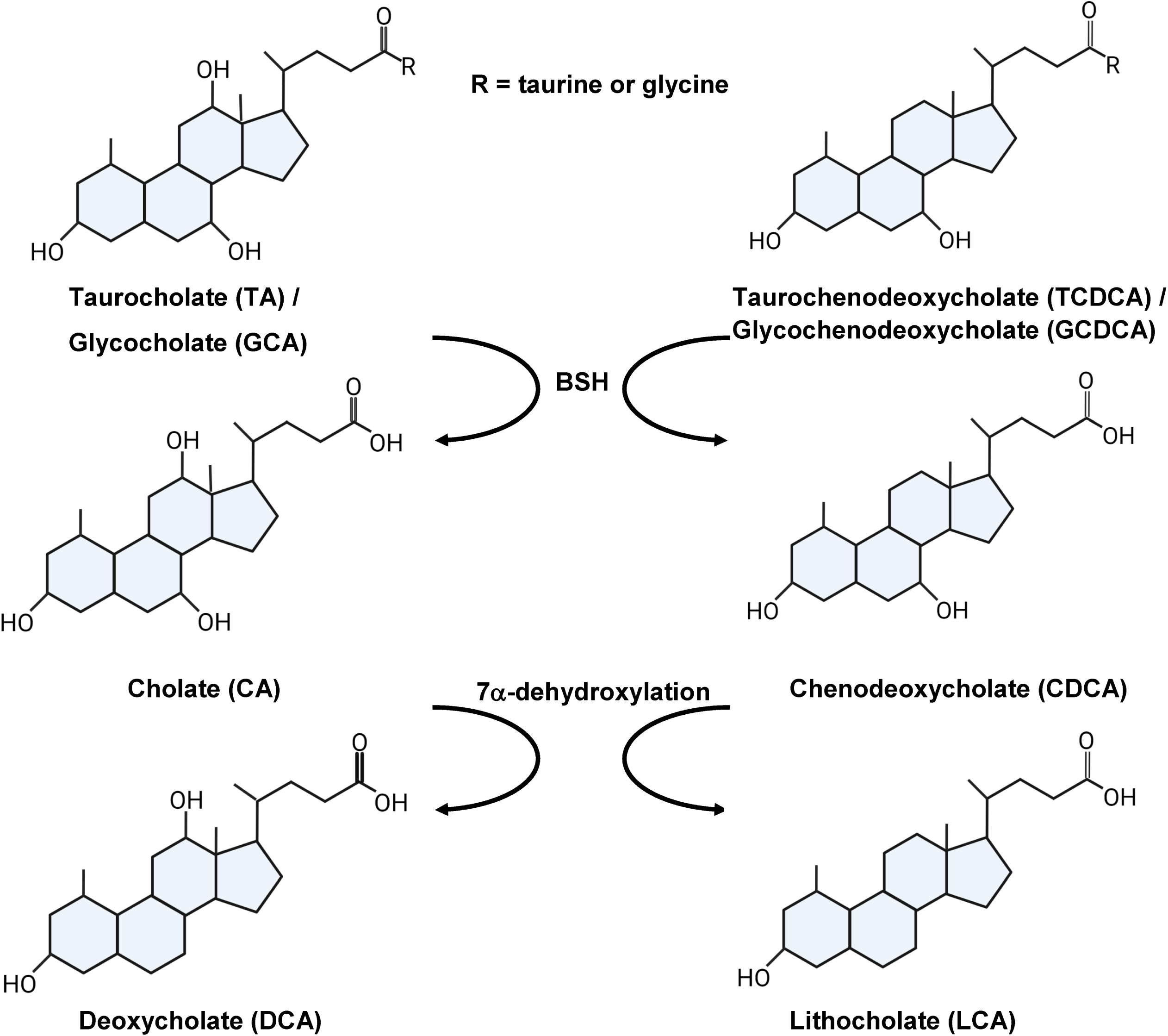
Host bile salts and derivatives. The two primary bile acids, cholate (CA) and chenodeoxycholate (CDCA), can be conjugated with taurine or glycine. Many bacteria in the gut can remove the conjugated amino acid (deconjugation) to generate the base bile salt. The deconjugated bile acids undergo 7α-dehydroxylation, performed by a small percentage of gut microbes, to generate the secondary bile salts deoxycholate (DCA) from CA and lithocholate (LCA) from CDCA.

Bile salt hydrolases (BSH) are part of the N-terminal nucleophilic (Ntn) hydrolase superfamily whose αββα-core structure is highly conserved (34). The presence of BSH genes is widely distributed. A study recently reported 69 bacterial genera in the HMP database that showed the presence of BSH genes, with the *Bacillus, Staphylococcus, Paenibacillus, Lysinibacillus, Clostridium*, and *Brevibacillus* genera having the greatest abundance of BSH genes (35). Although most bacteria encode only one BSH, there are some that encode up to four BSHs with the implication that multiple enzymes are expressed because of the well-established substrate specificity observed in BSH enzymes, thus allowing bacteria to deconjugate both taurine-conjugated and glycine-conjugated bile salts (31, 35, 36).

Bile salt hydrolase characterization in Clostridia is limited. Although the *C. perfringens* BSH enzyme is well-studied, with knowledge of the enzyme substrate specificity, cellular location and crystal structure (37-39), BSH enzymes in other Clostridia are less understood. This is despite apparent high abundance of BSH genes present, based on taxonomic and metagenomic data (31, 35, 40). To date, most experimental data on BSHs are derived from studies in Lactobacillus and Bifidobacterium species, that are commonly used for probiotics (32, 33, 35, 36, 41, 42).

To our knowledge, no prior studies have tested BSH activity in *C. difficile* strains. Understandably, *C. difficile* does not encode homologs of known BSH enzymes. However, herein, we show that several *C. difficile* strains, both laboratory- and epidemic-type strains, have BSH activity and that this activity is specific to taurine-conjugated bile salts. This activity is not restricted to vegetative cells as *C. difficile* spores also have a low amount of BSH activity, suggesting a potential role of the enzyme in *C. difficile* germination. Finally, we show that the identified BSH activity of *C. difficile* cells can generate a cholic acid source for *C. scindens. C. scindens* uses this CA source to generate DCA, thereby stimulating biofilm formation by *C. difficile* cells. Thus, our work shows that the identified BSH activity may provide *C. difficile* a mechanism to generate small amounts of CA for use by a competing microbe and stimulate its own biofilm formation and, thus, potential maintenance within a host.

## Results

### Different *C. difficile* ribotypes have substrate-specific bile salt hydrolase activity

In prior work, we tested the BSH of *C. scindens* and *C. hiranonis* and found that *C. scindens* cannot deconjugate bile salts but that *C. hiranonis* could (28). Surprisingly, when we included *C. difficile* as a supposed negative control, we found that the *C. difficile* CD630Δerm strain could remove the taurine from TA to generate CA. This was observed by incubating *C. difficile* vegetative cells overnight in rich medium supplemented with 1 mM TA. Subsequently, the bile salts were extracted, separated by HPLC and detected by evaporative light scattering (Figure S1). As shown in Figure 2A, *C. difficile* CD630Δerm could produce CA during this incubation; hyodeoxycholate (HCA) was included as an internal standard.

**Figure 2.**
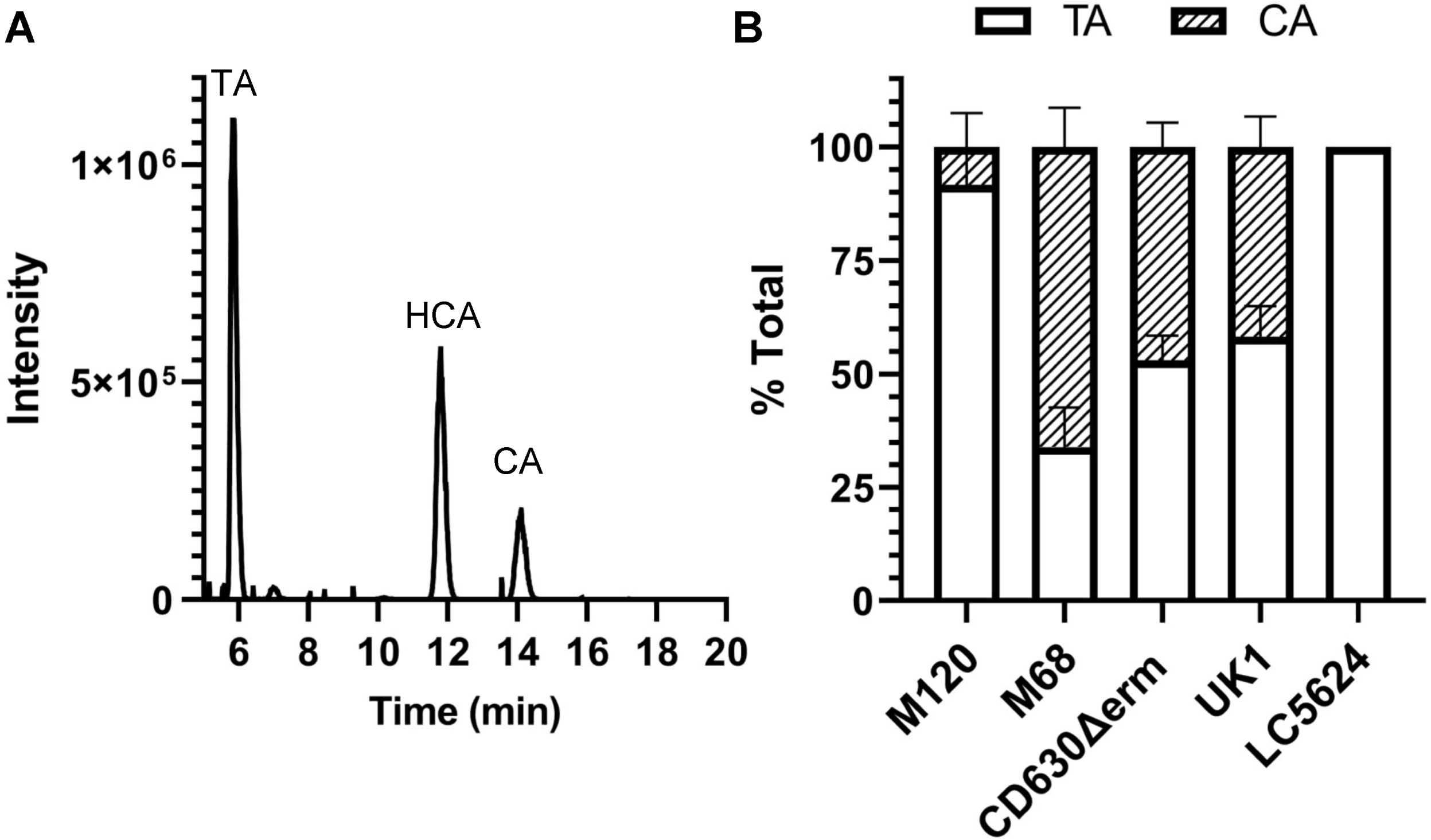
Several *C. difficile* strains can deconjugate taurocholate. A) *C. difficile* CD630Δerm was grown in rich medium supplemented with 1 mM taurocholate (TA) for 24 hrs. The bile salts were then analyzed by HPLC-coupled evaporative light scattering. Peak retention times for each bile acid were determined by using bile acid standards (Figure S1). B) Deconjugation of TA by *C. difficile* strains M120, M68, CD630Δerm, UK1 and LC5624 were determined as in (A). Values were calculated by measuring the peak areas for TA and CA and expressed as a percentage of total input. HDCA was included as an internal standard. Values shown are the average of three different independent experiments and error bars represent the standard error of the mean.

To understand if other *C. difficile* strains also had BSH activity, we grew strains of *C. difficile* derived from different ribotypes in the presence of taurocholate and quantified the amount of deconjugation. Surprisingly, in addition to *C. difficile* CD630Δerm (ribotype 014), we observed deconjugation of TA in 3 other strains [*C. difficile* M120 (ribotype 078), *C. difficile* M68 (ribotype 017), *C. difficile* UK1 (ribotype 027)] (Figure 2B). The *C. difficile* LC5624 strain (ribotype 106) did not deconjugate TA (Figure 2B). However, *C. difficile* M68 deconjugated approximately 70% of the TA to CA (Figure 2B). *C. difficile* CD630Δerm and UK1 deconjugated approximately 50% of the TA to CA and *C. difficile* M120 had the poorest activity with only ∼10% (Figure 2B).

To understand if this activity was specific to TA or if other taurine-derive bile salts could be deconjugated, we tested taurodeoxycholate (TDA) and taurochenodeoxycholate (TCDCA). The TDCA and TCDCA bile salts were deconjugated by all five *C. difficile* strains tested, including *C. difficile* LC5624 (Figure 3A and 3B). Similar to what we observed for deconjugation of TA, the levels of deconjugation for TDCA and TCDCA varied between strains. TCDCA showed the lowest deconjugation levels from the three tested taurine bile salts (Figure 3B) and the *C. difficile* LC5624 strain, ribotype 106 and the predominant ribotype causing CDI infections in the US at present (43), showed the lowest levels of BSH activity (Figure 3).

**Figure 3.**
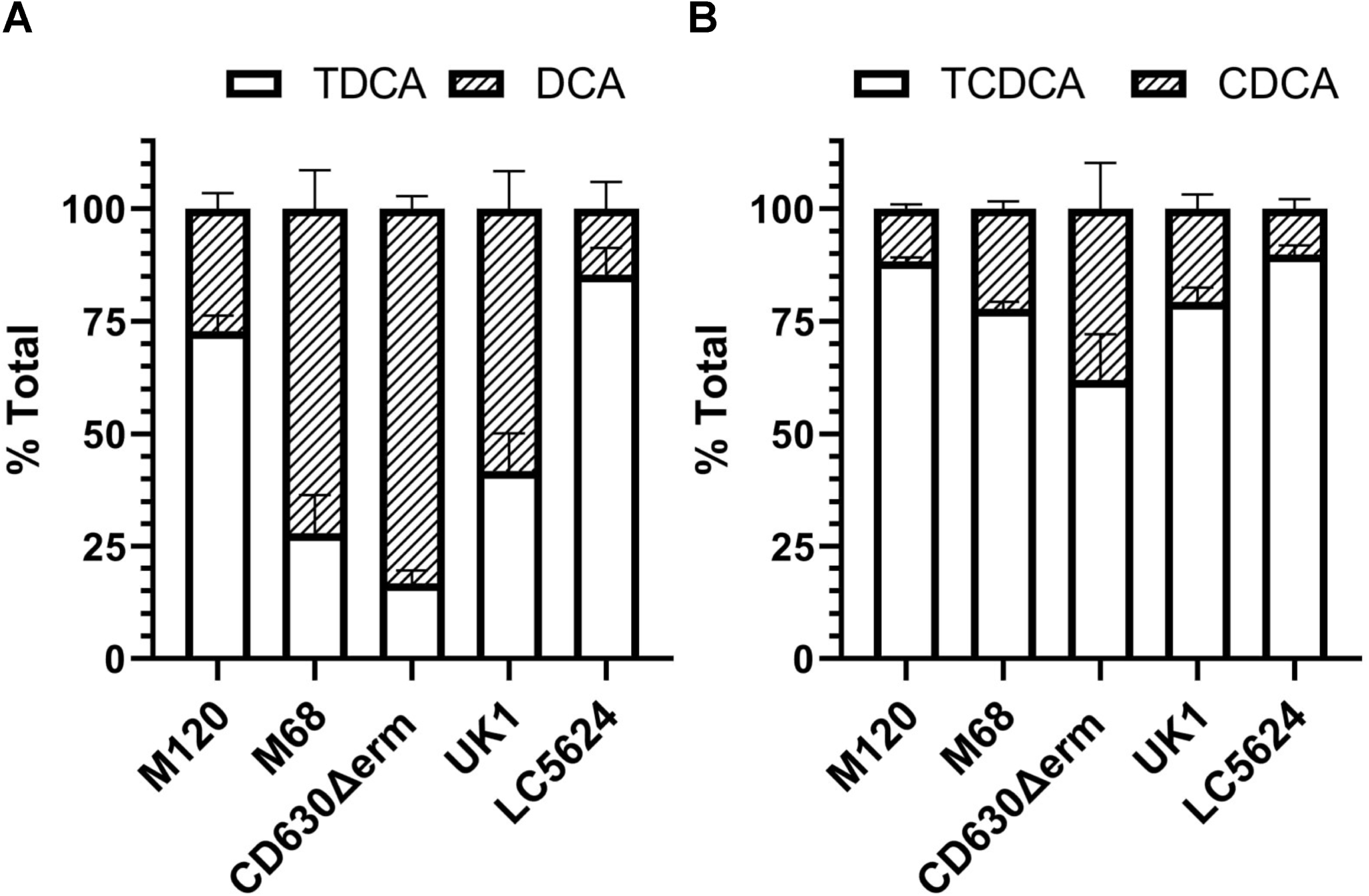
*C. difficile* deconjugates taurodeoxycholate and taurochenodeoxycholate. Deconjugation of taurodeoxycholate (TDCA) (A) or taurochenodeoxycholate (TCDCA) (B) by *C. difficile* strains M120, M68, CD630Δerm, UK1 and LC5624 were determined as in Figure 2. Values were calculated by measuring the peak areas for TDCA and DCA (A) or TCDCA and CDCA (B) and expressed as a percentage of total input. Hyodeoxycholate (HDCA) was included as an internal standard. Values shown are the average of three different independent experiments and error bars represent the standard error of the mean.

### *C. difficile* BSH activity is specific to taurine-derived bile salts

Because many BSH enzymes have specificity to one conjugated form or another, we next tested if *C. difficile* BSH activity is restricted to either the taurine-derived bile salts or if *C. difficile* can also deconjugate glycine-conjugated bile salts. *C. difficile* strains were grown in rich medium supplemented with a glycine-conjugated bile salt and analyzed as described above. *C. difficile* CD630Δerm could not deconjugate GCA (Figure 4A) or GCDCA (Figure 4B). Moreover, none of the tested strains had any BSH activity against the GCA (Figure 4C) or GCDCA (Figure 4D). Taken together, our data show the presence of BSH activity that is conserved throughout several *C. difficile* ribotypes and that this BSH activity is taurine-specific.

**Figure 4.**
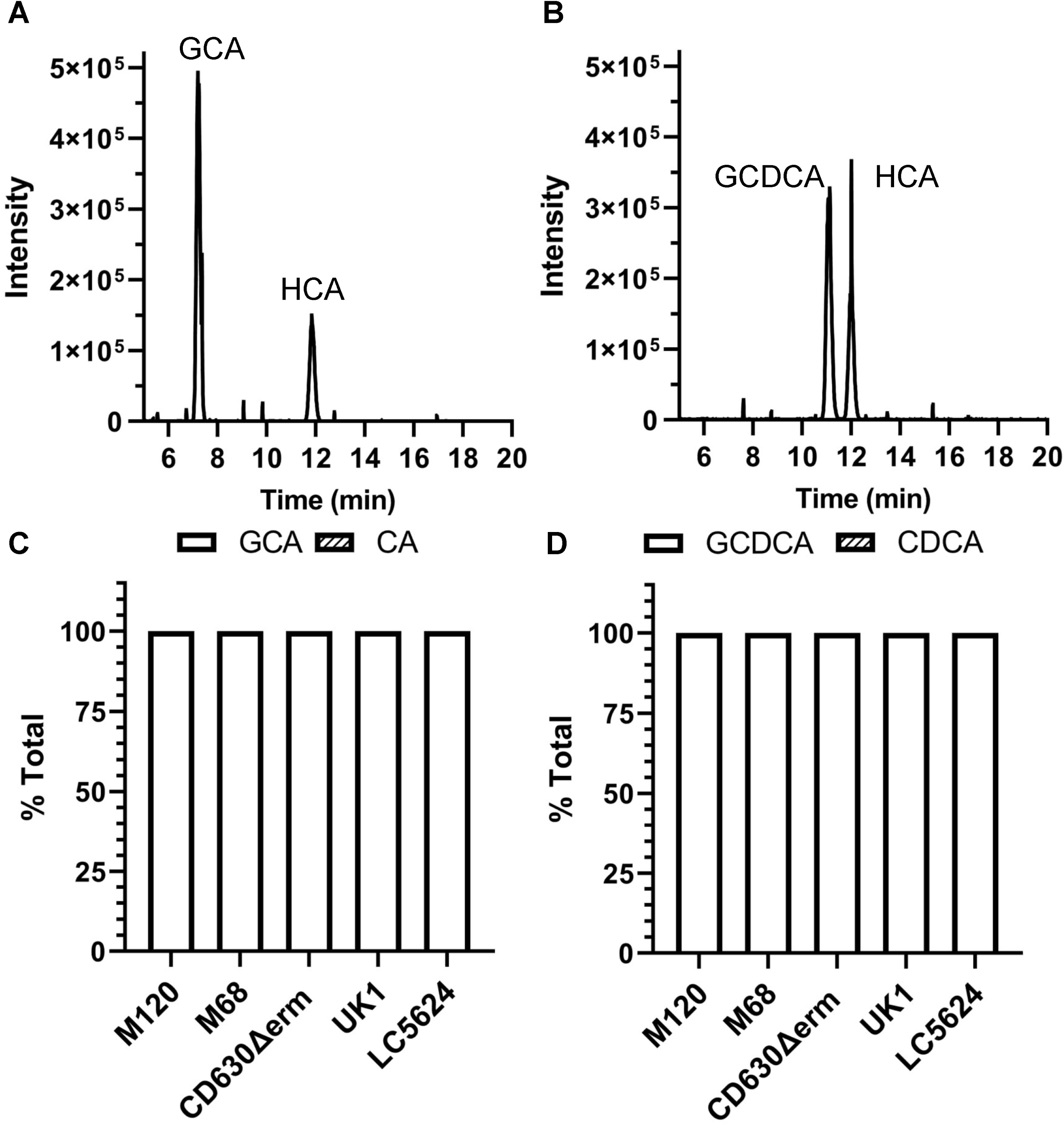
*C. difficile* BSH activity is taurine-specific. *C. difficile* CD630Δerm was grown in rich medium supplemented with 1 mM glycocholate (GCA) (A) or glycochenodeoxycholate (GCDCA) (B) for 24 hrs. The bile salts were then analyzed HPLC-coupled evaporative light scattering. Peak retention times for each bile acid were determined by using bile acid standards (Figure S1). Deconjugation of GCA (C) or GCDCA (D) by *C. difficile* strains M120, M68, CD630Δerm, UK1 and LC5624 were determined as in (A & B). Values were calculated by measuring the peak areas for GCA and CA or GCDCA and CDCA and expressed as a percentage of total input. HDCA was included as an internal standard. Values shown are the average of three different independent experiments and error bars represent the standard error of the mean.

### Dormant *C. difficile* spores deconjugate taurine derived bile salts

It is well established that the most effective germinants for *C. difficile* spores are TA and glycine (5, 6, 18). Because we observed BSH activity in *C. difficile* vegetative cells, we next tested if *C. difficile* spores also have BSH activity. To test for presence of BSH activity in dormant *C. difficile* spores, we purified spores and incubated them in buffer supplemented with 1 mM bile salts but no cogerminants (*e*.*g*., glycine; to prevent the spores from germinating). Though we observed BSH activity in spores, it was much lower when compared to the activity observed in vegetative cells (Figure 5). However, we observed that spores deconjugated: TA (Figure 5A), TDCA (Figure 5B) and TCDCA (Figure 5C).

**Figure 5.**
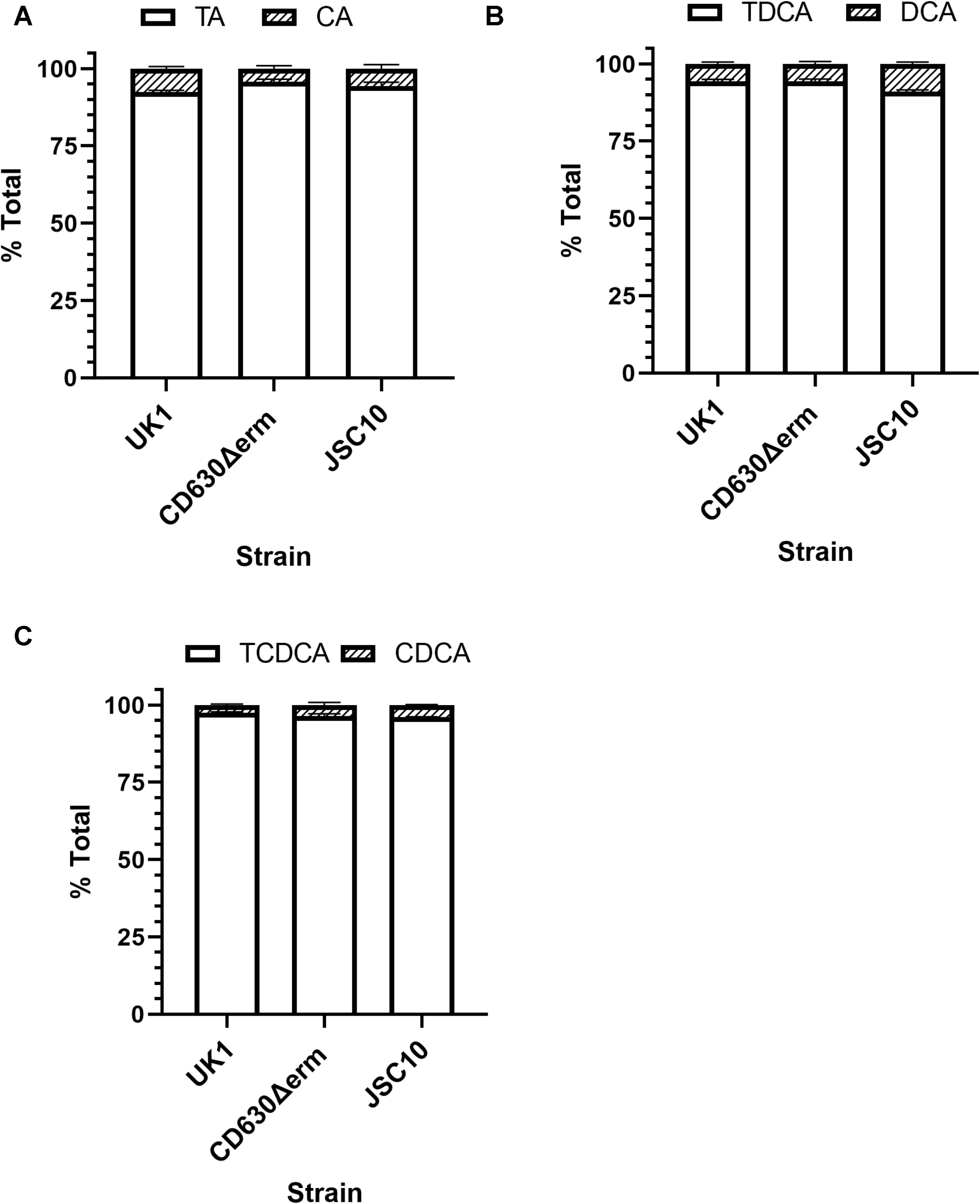
*C. difficile* spores show low levels of BSH activity towards taurine-derived bile acids. 1.2 × 10^9^ spores were incubated for 24 hours in an aerobic environment at room temperature in phosphate buffer saline (PBS) supplemented with A) 1mM TA, B) 1mM TDCA and C) 1 mM TCDCA. Samples were centrifuged and filter sterilized. Bile salts were measured using HPLC. Values were calculated by measuring the peak areas for the indicated bile salt, and expressed as a percentage of total input. HDCA was included as an internal standard. Values shown are the average of three different independent experiments and error bars represent the standard error of the mean.

To test if the spore-derived BSH activity was a result of low levels of autogermination of spores to the vegetative form, which could then provide BSH activity, we tested BSH activity in spores derived from the *C. difficile* JSC10 strain. *C. difficile* JSC10 has a mutation in the gene coding for the bile salt germinant receptor, CspC, and as such does not germinate in response to bile salts (8). As we observed for spores derived from the wildtype strains, the *C. difficile* JSC10 strain also had BSH activity (Figure 5A-5C).

### *C. difficile* can feed CA to *C. scindens* for DCA generation

Bile salt-metabolizing bacteria (*e*.*g*., *C. scindens* or *C. hiranonis*) are well-documented to correlate with a colonic environment that resists *C. difficile* colonization (26, 44, 45). Though the mechanism by which these bacteria protect again CDI may be through metabolic competition (28, 46-49), *C. difficile* must interact with these bile salt biotransforming bacteria. We decided to analyze potential coupling functions between *C. difficile* and other Clostridial species present in the GI tract. As we showed previously, *C. scindens* VPI12708 did not have BSH activity against TA, TDCA or TCDCA (Figure 6A). However, when grown in the presence of CA, *C. scindens* 7α-dehydroxylated CA to generate DCA (Figure 6A) (we did not observe generation of LCA from CDCA in our *C. scindens* strain). When *C. hiranonis* was grown in the presence of TA, it was able to efficiently deconjugate TA to CA and deconjugated TDCA and TCDCA [Figure 6B; (28)]. However, unlike *C. scindens*, our *C. hiranonis* 10542 strain did not 7α-dehyroxylate CA to generate DCA (Figure 6B).

**Figure 6.**
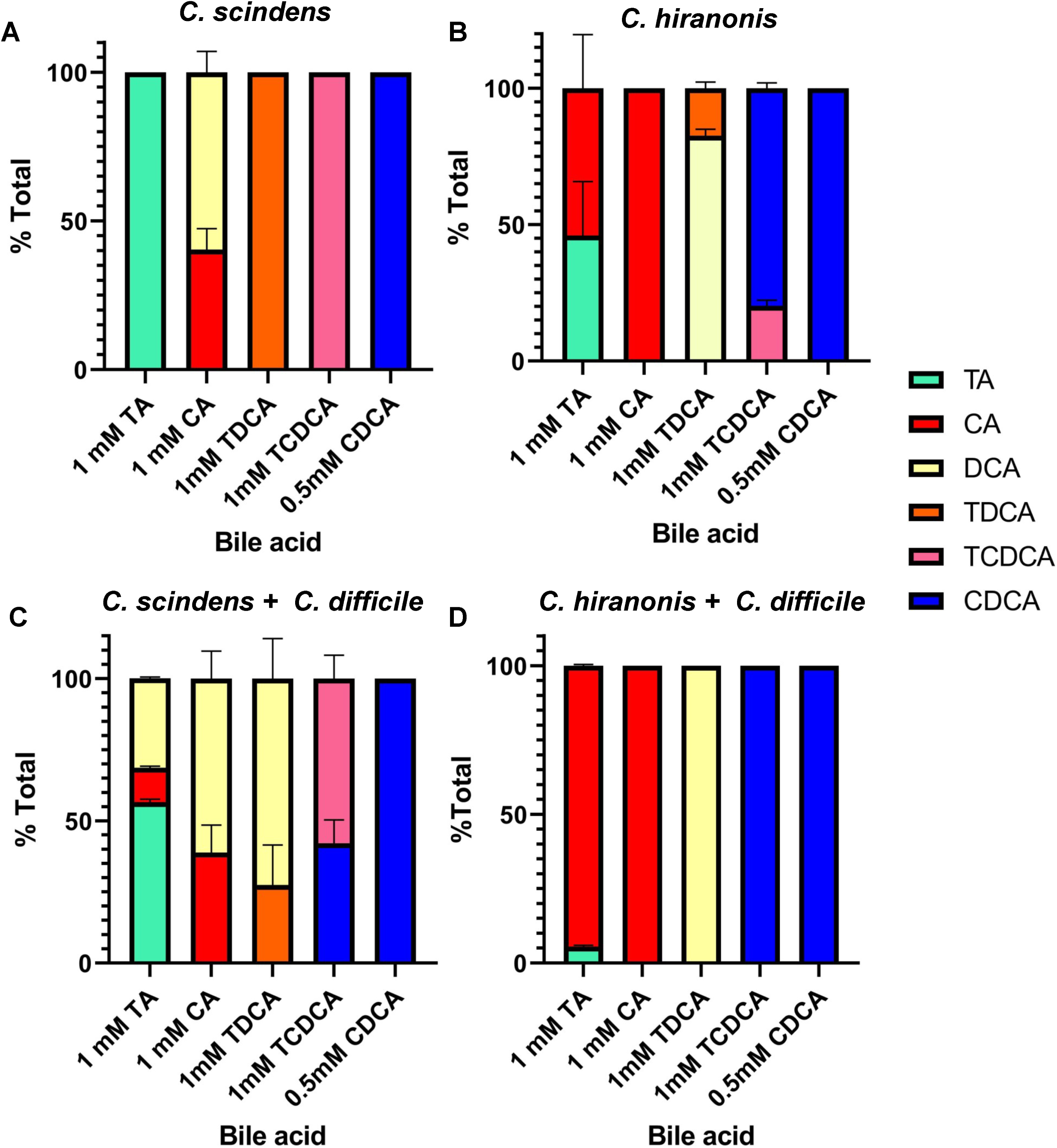
*C. difficile* BSH activity can feed CA to gut microbes. *C. scindens* (A) or *C. hiranonis* (B) overnight cultures were back diluted to 10^8^ CFU / mL and grown in presence of the indicated bile salts for 24 hrs at 37 °C in an anaerobic environment. C) *C. scindens* and *C. difficile* or D) *C. hiranonis* and *C. difficile* overnight cultures were back diluted to 10^6^ CFU / mL and grown in the presence of the indicated bile salts for 24 hrs at 37 °C in an anaerobic environment. Values were calculated by measuring the peak areas for the indicated bile salt and expressed as a percentage of total input. HDCA was included as an internal standard. Values shown are the average of three different independent experiments and error bars represent the standard error of the mean.

We next co-cultured *C. difficile* and *C. scindens* in the presence of different bile salts. We hypothesized that if *C. difficile* can deconjugate TA to CA, that *C. scindens* could use this CA to produce DCA. Indeed, when *C. difficile* and *C. scindens* were grown in rich medium supplemented with TA, we observed DCA production (Figure 6C). Similar to when *C. difficile* was grown in the presence of TDCA and TCDCA, we still observed deconjugation of these molecules (Figure 6C) but not the conversion of CDCA to LCA, similar to what we observed when *C. scindens* was grown in isolation (Figure 6A). When *C. difficile* and *C. hiranonis* were co-cultured, the amount of TA deconjugated was greater than when either of these bacteria were grown alone (Figure 2, Figure 6B). However, and as expected because *C. hiranonis* did not generate DCA from CA when grown in isolation (Figure 6B), we also did not observe DCA production in co-culture conditions. These results suggest *C. scindens* and *C. difficile* could interact and that bile salts could mediate this interaction.

### *C. difficile* can promote its own biofilm formation using BSH activity

In prior work, Dubois and colleagues found that 240 μM DCA promotes biofilm formation in *C. difficile* CD630Δerm (50). Because *C. difficile* BSH activity can produce CA from TA, we hypothesized that *C. difficile* may indirectly stimulate its own biofilm formation by feeding CA to *C. scindens*. To test this hypothesis, biofilm assays were run in an anaerobic chamber for 72 hr in the presence of TA or CA or DCA, as described previously (51). Subsequently, bile salt concentrations were determined and biofilms quantified using crystal violet staining. During co-culture of *C. difficile* and *C. scindens*, growth in the presence of TA yielded robust biofilm formation (Figure 7A). As expected, and as a positive control for biofilm formation, growth in the presence of DCA yielded robust biofilm (Figure 7A). However, we could not recapitulate biofilm levels observed in CA presence, despite the production of DCA in these conditions (Figure 7B). Because our *C. hiranonis* strain either does not generate DCA or does so below the limit of detection for our instrument (0.2 nmol), we tested if merely growing *C. difficile* in the presence of another metabolic competitor (28) induces biofilm formation by *C. difficile* cells (Figure 7C). Despite co-culturing *C. difficile* and *C. hiranonis* in the presence of either TA or CA, we did not observe biofilm formation (Figure 7C). However, growth in the presence of DCA, as expected, generated robust biofilm (Figure 7C).

**Figure 7.**
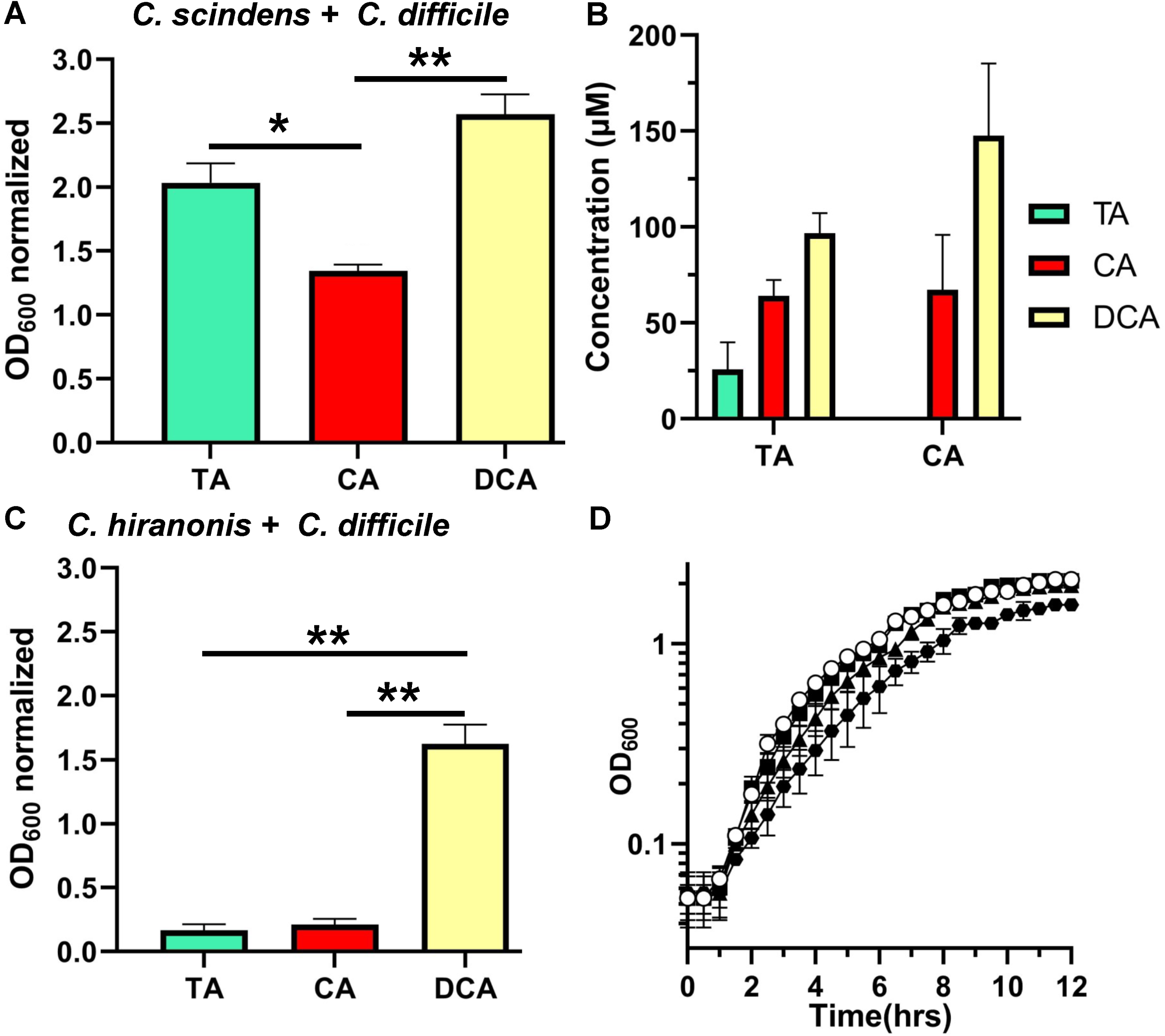
*C. difficile* BSH activity contributes to biofilm formation in a *C. difficile*- *C. scindens* dual species biofilm. A) *C. difficile* was co-cultured with *C. scindens* and the indicated bile salt and the amount of biofilm formation was quantified using a crystal violet assay. B) Concentrations of the indicated bile salts in the biofilm assays. C) *C. difficile* was co-cultured with *C. hiranonis* and the indicated bile salt and the amount of biofilm formation was quantified using a crystal violet assay. D) Overnight cultures of *C. difficile* CD630Δerm were back diluted to an OD_600_ 0.05 and monitored for 12 hours in rich medium alone ((○) or supplemented with 100 µM deoxycholate (DCA) (■), 150 µM DCA ((▲), 240 µM DCA ((●). Values shown are the average of three different independent experiments and error bars represent the standard error of the mean. Statistical analysis was performed using an ordinary one-way ANOVA. * p < 0.005, ** p < 0.001.

We next quantified the amount of bile salt present in the biofilm conditions. When *C. difficile* and *C. scindens* were co-cultured in the presence of TA, approximately 75 μM CA was generated and approximately 100 μM DCA was generated (Figure 7B). When grown with CA, approximately 150 μM DCA was produced. Secondary bile salts are well-documented to inhibit *C. difficile* growth. To understand if the DCA produced by *C. scindens* can inhibit *C. difficile* growth, we grew *C. difficile* in 100 μM, 150 μM, or 240 μM DCA. As shown in Figure 7D, these concentrations of DCA did not significantly inhibit *C. difficile* growth, though there was a trend of less growth at 240 μM DCA. These results indicate that the amount of DCA produced by *C. scindens* in response to deconjugation of TA by *C. difficile* does not inhibit growth but, that *C. difficile* responds to this by producing biofilm.

## Discussion

BSH research has increased significantly since their discovery in the 1970’s due to their potential to modulate bile salts in the gut environment, the health implications in bile salt modulation, as well as bile salt toxicity towards gut microbiota (32, 33, 36, 41, 52). Nevertheless, most BSH research has focused on bacterial species that are used as probiotics, whereas studies aiming at characterization and function of BSH in human pathogens is limited. Here, we provided initial characterization of the BSH activity in the human pathogen *C. difficile*. Our results show that BSH activity is conserved across ribotypes and that deconjugation is specific to taurine-conjugated bile salts (Figures 2 and 3). *C. difficile* does not encode orthologues of known BSH genes. Based upon recent work (33, 36), we attempted to bioinformatically determine if *C. difficile* had a weak homolog but this proved fruitless. At its basis, BSH activity is the cleavage of an amide bond. We hypothesize that one of the many amidases or peptidases encoded by *C. difficile* either is a novel BSH or moonlights as a BSH. We are continuing our efforts in identifying this enzyme.

It is unclear why *C. difficile* BSH activity favors taurine-conjugated bile salts. One hypothesis is the use of taurine for Stickland fermentation. However, glycine is only one of three amino acids that can be used in the reductive branch of *C. difficile* Stickland metabolism. Thus, the ability to use glycine released by glycine-derived bile salts should also be beneficial to *C. difficile* (46, 53-55). On the other hand, recent discoveries of bile salts as modulators of *C. difficile* TcdB effectiveness showed the ability of the taurine conjugated bile salt TCDCA, to inhibit toxin activity in a dose dependent manner (56). The BSH activity by different *C. difficile* isolates to deconjugate TCDCA (Figure 3B) and other taurine bile salt derivatives may be a mechanism by the bacterium to modulate toxin activity depending on the surrounding conditions.

Germination by *C. difficile* spores is well-established to be influenced by and dependent on host bile salts (5, 7-9, 11, 14-19, 57). Although all cholic acid-derived primary bile salts promote germination (5), a study testing the germination of individual *C. difficile* spores found that TA and TDCA had the highest germination levels after one hour (58). Additionally, although most chenodeoxycholic acid-derived bile salts inhibit germination, TCDCA does not inhibit spore germination (data not shown). Because we observe low BSH activity in *C. difficile* spores, we hypothesize that the observed activity is due to the misincorporation of the enzyme responsible for this activity into the coat layer of the developing spore. Still, this could provide a means to modulate the efficacy of germination by converting TCDCA to CDCA, a competitive inhibitor of spore germination. This sort of mechanism is observed for germination in bacilli where L-alanine stimulates spore germination but a spore-associated alanine racemase converts L-alanine to D-alanine, an inhibitor of spore germination (59-62).

During growth in a host, *C. difficile* encounters an environment that has a diverse repertoire of bile salts. The ability to use this information to affect changes to the community would provide an advantage for *C. difficile* in the host. Previously, we hypothesized that *C. difficile* competes with *C. scindens*, and related bacteria, for proline and / or glycine (28). During a successful infection, *C. difficile* also consumes proline and glycine which may exclude *C. scindens* from regaining a foothold to provide colonization resistance. However, should *C. scindens* begin to accumulate in the GI, this could signal to *C. difficile* that an environment that is non-conducive for optimum growth is being established. To resist these changes and maintain itself, formation of a biofilm would be advantageous. We hypothesize that *C. difficile*-mediated deconjugation of TA results in the environment being ‘seeded’ with CA. As *C. scindens* becomes abundant in the GI, *C. scindens* converts the generated CA to DCA. The generated DCA signals back to *C. difficile* to initiate biofilm production and maintenance in the host (Figure 8). Moreover, *C. difficile* could take advantage of high TA levels during the initial stages of infection not only for germination (63), but for generation of CA that can be used by other bacteria that do not encode BSH enzymes to generate DCA and persist in the gut environment and, potentially, to cause recurring infections.

**Figure 8.**
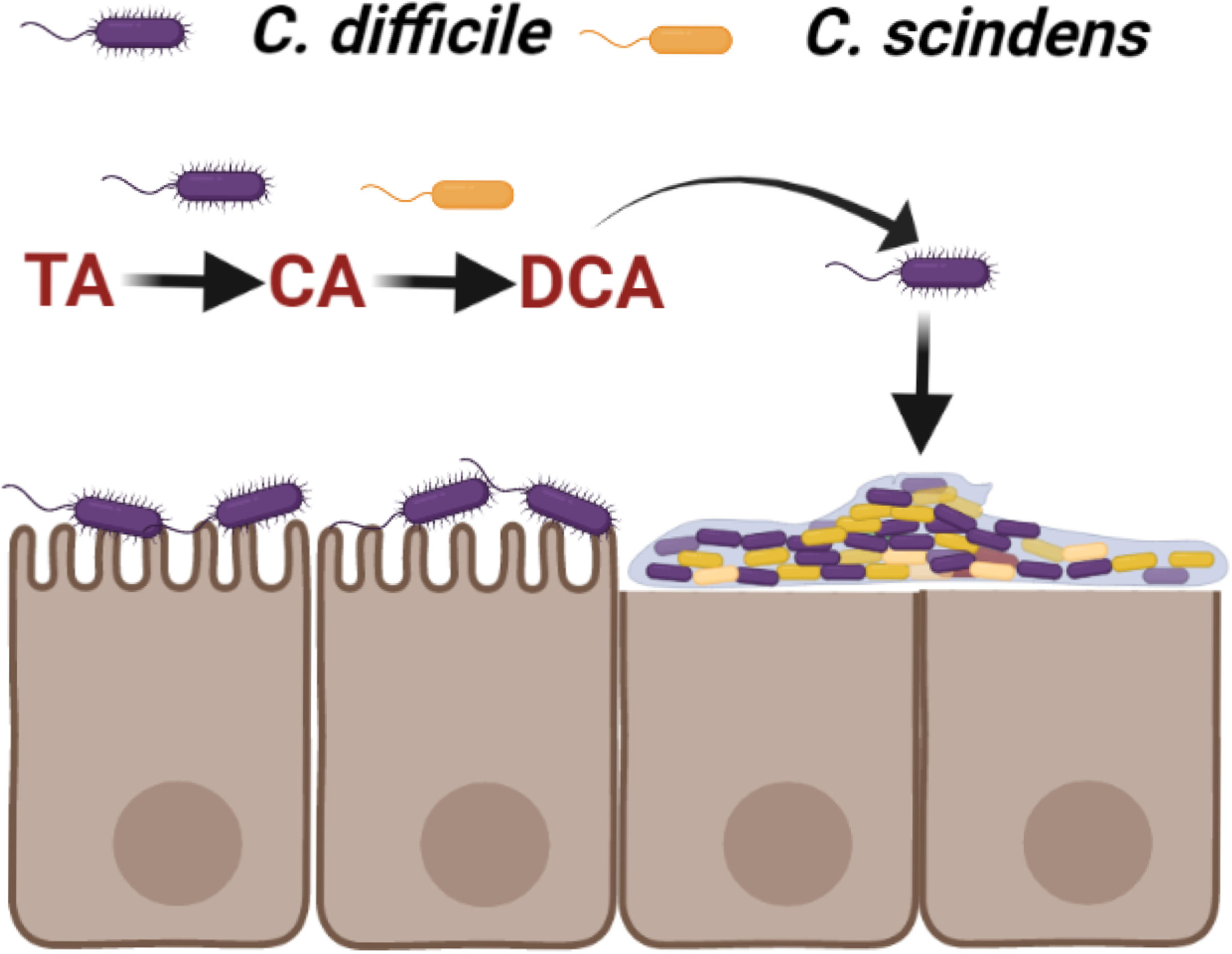
Working model for DCA-mediated dual species biofilm formation. The ability of *C. difficile* to deconjugate TA during colonization “seeds” the gut environment with CA to be used by *C. scindens* during gut microbiome recovery. The presence of DCA signals a potentially non-conducive environment for *C. difficile* growth and the bacteria forms a DCA-mediated biofilm to persist in the gut.

## Materials and Methods

### Bacterial and strains

Clostridial strains, were grown at 37 °C in an anaerobic chamber (Coy Laboratories; model B; >4% H_2_, 5% CO_2_, 85% N_2_) on brain heart infusion agar (BHI) and 0.1% L-cysteine, BHI supplemented with 5 g / liter yeast extract and 0.1% L-cysteine (BHIS) or BHIS with 100 mM glucose and 0.1% L-cysteine (BHISG). Spores were generated on 70:30 agar medium [63 g / L Bacto peptone, 3.5 g / L protease peptone, 11.1 g / L BHI, 1.5 g / L yeast extract, 1.06 g / L tris base and 0.7 g / L ammonium sulfate (NH_4_SO_4_)].

### BSH single and dual species activity assays

Clostridial species were grown on BHI medium with 0.1 % L-cysteine for 16 hours. Cultures were back diluted to 10^8^ CFU into fresh BHI medium supplemented with 1 mM / 0.5 mM of specified bile salt. For dual species assays, overnight cultures were back diluted to 10^6^ CFU, and added to BHI medium, in a 1:1 ratio, with specified bile salts. Cultures were grown for 24 hours and then centrifuged for 10 minutes at 4,000 x g. The pellet was suspended in 100% methanol and the supernatant was lyophilized and resuspended in already suspended pellet solution. The presence of specific bile salts in samples was measured as described below.

### Bile salt separation

Bile salts were separated by reverse-phase HPLC using a Shimadzu prominence HPLC system. Twenty-five microliter samples were separated using a Synchronis C18 column (4.6 by 250 mm; 5 μm particle size; ThermoFisher 97105–254630) using a mobile phase consisting of 53% methanol, 24% acetonitrile, 23% water and 30 mM ammonium acetate (pH 5.6). Bile salt peaks were detected using a Sedere Sedex model 80 LT-ELSD (low temperature-evaporative light scattering detector) using an air pressure of 50 psi of Zero Grade air at 94 °C. Different amounts of specific bile salts [taurocholic acid (Sigma Aldrich 86339-25G), glycocholic acid (Sigma Aldrich G7132-1G), taurochenodeoxycholic acid (Sigma Aldrich T6260-250MG), glycochenodeoxycholic acid (Sigma Aldrich G0759-500MG), hyodeoxycholic acid (Sigma Aldrich H3878-5G), chenodeoxycholic acid (Acros organics C9377-25G), cholic acid (Sigma Aldrich C1129-100G), deoxycholic acid (Sigma Aldrich D2510-100G), lithocholic acid (Acros organics L6250-5G), taurodeoxycholate (Sigma Aldrich T0875-5G)] were separated to generate standard curves. The area under each peak was calculated and plotted against the concentration of bile salt added and a trend line was generated for each bile salt. Concentration of the bile salts in samples (nmol) was calculated using the standard curves of pure bile salts and normalized with the added internal standard (HDCA). Percent total was calculated by dividing concentration of specific bile salt over total bile salts presence.

### *C. difficile* spores BSH assays

Spores of *C. difficile* strains were purified as described below on 70:30 sporulation media. Following purification, spores were counted and 1.5 ×10^8^ spores were used for each BSH assay. Purified spores were incubated in PBS supplemented with 1mM TA, TDCA, or TCDCA fer at room temperature for 24 hours in an aerobic environment. The presence of bile salts was measured by HPLC-ELSD as described above.

### Spore purification

Spores were purified as previously described (10, 16, 36, 65). Briefly, strains were grown on 70:30 sporulation medium. After 5 days, growth from 7 plates each were scraped into 1 mL distilled water (dH_2_O) in microcentrifuge tubes and left overnight at 4 °C. The cultures were then resuspended on dH_2_O in the same microcentrifuge tubes and centrifuged at >14,000 × g for 1 min, the top layer containing vegetative cells and cell debris was removed by pipetting, and the rest of the sediment resuspended in fresh dH_2_O. The tubes, again, were centrifuged for 1 min at >14,000 × g, the top layer removed, and the sediment resuspended. This was repeated 5 more times, combining the sediment from 7 tubes into one. The spores were then separated from the cell debris by centrifugation through 50% sucrose for 20 min at 4°C and 3,500 × g. The resulting spore pellet was then washed 5 times with dH_2_O, resuspended in 1 mL dH_2_O, and stored at 4 °C until use.

### Dual species biofilm assays

Biofilms assays were done as previously reported with some modifications. Briefly, overnight cultures of *C. difficile, C. scindens* and *C. hiranonis* in BHIS were normalized to an OD_600_ nm of 1.00 and an equal volume of each (10 µL) was added to prepared wells (final volume 1 mL: 24-well plate). Filter sterilized bile salts were introduced into equilibrated BHISG medium at 190 µM - 240 µM. Cultures were incubated in 24-well tissue culture treated plates and the plates were incubated for 72 h at 37 °C in an anaerobic chamber. After 72hr, plates were taken out of anaerobic environment and spent media was removed by washing individual wells twice with phosphate-buffered saline (PBS). Biofilms were air dried and stained with crystal violet (CV; 0.2% w/v) for 20 min. CV was removed by inversion; wells were washed twice with PBS then air-dried.

Dye bound to the biofilm biomass was solubilized by adding 1 mL of a 75% ethanol solution and the absorbance, corresponding to the biofilm biomass, was measured at 600 nm with a plate reader (Promega GloMax Explorer). When necessary, the solubilized dye was diluted for the reading to remain in the linear range of the spectrophotometer. Because of individual variations in negative control wells between rows, each row in the 24-well plates was normalized to their row negative control. *C. difficile* - *C. scindens* dual species biofilm was normalized to the levels of biofilm found in a *C. scindens*-only control. Unwashed biofilms were used to measure presence of primary and secondary bile salts using the above described HPLC-ELSD method.

### Growth curves

An overnight culture of *C. difficile* CD630Δerm in BHISG was back diluted to an OD_600_ 0.05 into fresh BHISG alone or in BHISG supplemented with 150 µM, 200 µM, or 240 µM DCA. OD_600_ was recorded every half hour for a total of twelve hours.

### Statistical analyses

Data represent results from at least three independent experiments, and the error bars represent standard errors of the means. One-way ANOVA analysis was performed using GraphPad Prism version 9.0.2 (161) for Windows (GraphPad Software, San Diego, California USA).

## Acknowledgements

This project was supported by awards R01AI116895, U01AI124290 and R21AI144454 from the National Institute of Allergy and Infectious Diseases. Andrea Martinez Aguirre is a recipient of a CONACYT-COECYT fellowship 2017-2022 scholar/scholarship 625561/472087. The content is solely the responsibility of the authors and does not necessarily represent the official views of the NIAID. The funders had no role in study design, data collection and interpretation, or the decision to submit the work for publication.

---

**Figure S1. Bile acid standards using HPLC** Indicated bile acids were separated by High-Performance Liquid Chromatography. Compounds were detected by evaporative light scattering. Standard curves were generated by calculating the area under the corresponding peaks.

